# Elevation of *Clavibacter michiganensis* subsp. *californiensis* to species level as *Clavibacter californiensis* sp. nov., merging and re-classification of *Clavibacter michiganensis* subsp. *chilensis* and *Clavibacter michiganensis* subsp. *phaseoli* as *Clavibacter chilensis* sp. nov. based on complete genome *in-silico* analyses

**DOI:** 10.1101/2022.01.17.476658

**Authors:** Dario Arizala, Shefali Dobhal, Anne M. Alvarez, Mohammad Arif

## Abstract

The Gram-positive *Clavibacter* genus is currently divided into seven species (*C. michiganensis*, *C. nebraskensis*, *C. capsici*, *C. sepedonicus*, *C. tessellarius*, *C. insidiosus* and *C. zhangzhiyongii*) and three subspecies (*C. michiganensis* subsp. *californiensis*, *C. michiganensis* subsp. *chilensis* and *C. michiganensis* subsp. *phaseoli*). Recent studies have indicated that the taxonomic rank of the subspecies must be re-evaluated. In this research, we assessed the taxonomy position of the three *C. michiganensis* subspecies and clarified the taxonomic nomenclature of other 75 *Clavibacter* strains. The complete genomes of the type strains of the three *Clavibacter* subspecies, type strain of *C. tessellarius* and *C. nebraskensis* A6096 were sequenced using PacBio RSII technology. Application of whole-genome-based computational approaches such as average nucleotide identity (ANI), digital DNA-DNA hybridization (dDDH), multi-locus sequence analysis (MLSA) of seven housekeeping genes (*acnA, atpD, bipA, icdA, mtlD, recA* and *rpoB*), phylogenomic tree reconstructed from 1,028 core genes, and ANI-based phylogeny pinpointed conclusive evidence to raise *C. michiganensis* subsp. *californiensis* to the species status. These results led us to propose the establishment of *C. californiensis* sp. nov. as a species with its type strain C55ᵀ (=CFBP 8216ᵀ=ATCC BAA-2691ᵀ). Moreover, the orthologous and *in-silico* dot plot analyses, along with the aforementioned bioinformatic strategies, revealed a high degree of homology between *C. michiganensis* subsp. *chilensis* and *C. michiganensis* subsp. *phaseoli*. Based on these outcomes, we proposed to combine both subspecies into a single taxon and elevate its rank to the species level as *C. chilensis* sp. nov., with ZUM3936ᵀ (= ATCC BAA-2690ᵀ = CFBP 8217ᵀ) as the type strain.

## Introduction

The genus *Clavibacter*, described by Davis *et al*. [1] constitutes a set of Gram-positive bacterial phytopathogens of agricultural crops, namely, tomato, pepper, potato, wheat, corn and alfalfa [2]. Originally, the genus *Clavibacter* was solely composed by the species *C. michiganensis* [3], which was divided into seven pathogenic subspecies assigned according to its narrow host specificity including: *C. michiganensis* subsp. *michiganensis*, the causal agent of canker and wilt of tomato [3, 4]; *C. michiganensis* subsp. *sepedonicus*, responsible of potato ring rot [2]; *C. michiganensis* subsp. *insidiosus*, inducing wilting and stunting in alfalfa [5]; *C. michiganensis* subsp. *tessellarius*, pathogenic to wheat, producing leaf freckles and leaf spots in a mosaic pattern [6]; *C. michiganensis* susbp. *nebraskensis*, causing blight and wilting of maize [7]; *C. michiganensis* susbp. *capsici*, inducing canker in pepper [8] and *C. michiganensis* subsp. *phaseoli*, reported as the responsible species of bacterial bean leaf yellowing [9]. The first three aforementioned subspecies have been categorized in the A2 list as quarantine organisms by the European and Mediterranean Plant Protection Organization [10]. Additionally, two subspecies isolated from tomato seeds, and described as non-pathogenic were named as *C. michiganensis* subsp. *californiensis* and *C. michiganensis* subsp. *chilensis*, based on their geographic origin California and Chile, respectively [11]. In 2018, five of the pathogenic subspecies were elevated to species level and reclassified as *C. sepedonicus*, *C. insidiosus*, *C. tessellarius*, *C. nebraskensis* and *C. capsici* based on whole-genome and multilocus sequence analyses [12]. Later, this new taxonomy assignation was also supported by a genome-based taxonomic study conducted in the phylum *Actinobacteria* and the description of *C. michiganensis* subsp. *michiganensis*, the bacterial tomato pathogen, was emended as *C. michiganensis* [13]. Most currently, a new pathogenic bacterium causing leaf brown spot and decline of barley was described as *C. zhangzhiyongii* [14].

Accurate systematics as well as a concise and reliable taxonomy designation of prokaryotes are highly primordial steps in epidemiology, pathogen surveillance, genome-informed diagnostics and for an efficient disease management [15, 16]. Next generation sequencing has boosted the identification of unique genomic traits within a taxa and the development of novel whole-genome-based computational tools for bacterial taxonomy classification, namely: phylogenetic tree reconstruction using core-genome alignment [17] and species delineation through calculation of the Average Nucleotide Identity (ANI) [18, 19] and digital DNA-DNA hybridization (dDDH) [20, 21]. Inclusion of these approaches in phylogenomics have elucidated novel species, and helped to clarify the taxonomy position of complex and highly heterogeneous bacterial species, which would have not been possible by using merely former taxonomic procedures, such as phenotypic characterization, 16S ribosomal RNA sequencing and multi-locus sequencing typing/analysis (MLST/MLSA) [15, 16]. In *Clavibacter*, for instance, the use of high throughput taxonomic approaches allowed to reclassify some subspecies to species level [12], and to describe new species, namely *C. michiganensis* subsp. *californiensis* and subsp. *chilensis* [11]. However, when the above two subspecies were described, their respective genomes were not available in any database. In 2018, the draft genomes of these two isolates together with *C. michiganensis* subsp. *phaseoli* were published and available in the NCBI GenBank database [22]. Later, two comparative genomics studies, based on pairwise comparison analyses, showed ANI and dDDH values below the suggested 96- and 70-% threshold for species delineation, respectively [23, 24], pinpointing that the subspecies status of these three organisms has to be re-evaluated. Considering this background, the main objectives of the present research were: first, to assess and validate the taxonomy status of the three subspecies, *C. michiganensis* subsp. *californiensis*, *Clavibacter michiganensis* subsp. *chilensis* and *Clavibacter michiganensis* subsp. *phaseoli* by using high-quality complete genome sequences of their type strains, along with the complete genomes of the type strain of *C. tessellarius* and *C. nebraskensis* strain A6096; and second, to verify/correct the taxonomic description of other *Clavibacter* strains, whose assigned names in the NCBI database are incorrect, by using the modern whole-genome-based *in silico* tools for bacterial taxonomy description.

### Isolation, Ecology and Genome dataset preparation

In this study, a large panel of 80 *Clavibacter* strains belonging to different species and isolated through different years, geographic regions, hosts, and niches were used for the phylogenomics and whole-genome-based computational analyses. Among the 80 strains, the complete genomes sequences of the type strains of the three subspecies of *C. michiganensis* (subsp. *californiensis*, *chilensis* and *phaseoli*), along with the type strain of *C. tessellarius* and *C. nebraskensis* strain A6096 were used for the taxonomy assessments instead of their previous draft genome versions. The bacterial cultures of these five strains were obtained from the Pacific Bacterial Collection at the University of Hawai’i at Manoa, Honolulu, HI, United States. The isolates were acquired from −80°C and plated onto yeast sucrose calcium carbonate (YSC) medium (yeast extract 10 g/l, sucrose 20 g/l, calcium carbonate 20 g/l and agar 17 g/l) and incubated for 72 h at 26°C (±2°C). Complete or draft genomes of the other 75 *Clavibacter* strains were retrieved from the NCBI GenBank database as of October 13^th^, 2021. Detail descriptions of all strains used in this research are provided in Table S1.

### Phylogeny based on Multi-Locus Sequence Analysis (MLSA)

To determine the taxonomic position of the five *Clavibacter* genomes sequenced in this study and resolve the taxonomy status of the other 75 *Clavibacter* strains listed in Table S1, a multi-locus sequence analysis (MLSA) was performed using seven housekeeping genes *acnA, atpD, bipA, icdA, mtlD, recA* and *rpoB*. Multiple gene sequences were aligned using MUSCLE and then concatenated in Geneious Prime version 2021.1.1. The concatenated alignment was used to generate a maximum-likelihood (ML) phylogenetic tree based on a bootstrap test of 1,000 replicates. Phylogenetic evolutionary relationships were assessed in MEGA 11. All sequences of the seven housekeeping genes corresponding to the 75 *Clavibacter* strains were retrieved from complete or draft genomes available at the NCBI GenBank database (Table S1). *Rathayibacter iranicus* NCCPB2253 and *Rathayibacter toxicus* WAC3373 were used as outgroups for the phylogenetic analyses. On the phylogenetic tree (Fig 1), nine clear clusters were identified. The biggest and first cluster comprised 44 strains of the recently emended species *C. michiganensis* [13] and the second cluster grouped the type strain of the proposed species *C. californiensis* sp. nov. together with the strain AY1B2, isolated from ryegrass in 2013 and reported as *C. michiganensis* in the NCBI GenBank database. Following these two clusters, the third cluster harbored 2 sub-clades, the first composed of the strains VKM Ac-2886, isolated from *Sambucus racemosa*, and CFBP 7491, isolated from tomato seeds; the second sub-clade integrated the type strains of *C. michiganensis* subsp. c*hilensis* CFBP 8217ᵀ, isolated from tomato seeds, and *C. michiganensis* subsp. *phaseoli* LPPA 982ᵀ, pathogen of bean, highlighting that these two subspecies must be merged into a single species. We proposed the species name for these 4 strains within the third cluster as *C. chilensis* sp. nov. Clusters fourth to sixth constituted monophyletic clades comprising the species *C. nebraskensis*, *C. insidious* and *C. sepedonicus*. Our *C. nebraskensis* strain A6096 positioned close to the strains HF4 and CFBP 7577. Interestingly, the strain CFBP 7494, a tomato endophytic bacterium isolated from seeds in Chile, clustered closely to *C. insidiosus*, like previous studies [25], but in a different sub-clade. Clusters seventh and eight encompassed the pathogens of pepper (*C. capsici*) and barley (*C. zhangzhiyongii*), respectively. The endophytic strain CFBP 7576 submitted in the NCBI database as *C. michiganensis*, grouped with other *C. capsici* isolates, which is coherent with the fact that this bacterium was proved to cause disease on pepper [25]. The last cluster comprised the strains DOAB 609, CFBP 8017, the type strain of *C. tessellarius* and CFBP 3399. Intriguingly, CFBP 3399, named as *C. michiganensis* and isolated from tulip (*Tulipa* sp.) in Netherlands, formed its own clade with *C. tessellarius* type strain ATCC 33566ᵀ. We hypothesized that CFBP 3399 might be a pathogen of wheat similarly like the tomato endophyte CFBP 8017 that was observed to cause mosaic disease symptoms on wheat [25]. Although DOAB 609 and CFBP 8017 are closely related to ATCC 33566ᵀ, former analysis based on genome-to-genome similarity indices has shown that these strains should not be classified as *C. tessellarius* but as new species [23]. Indeed, our analyses, described further, match with these previous assumptions.

**Figure 1.**
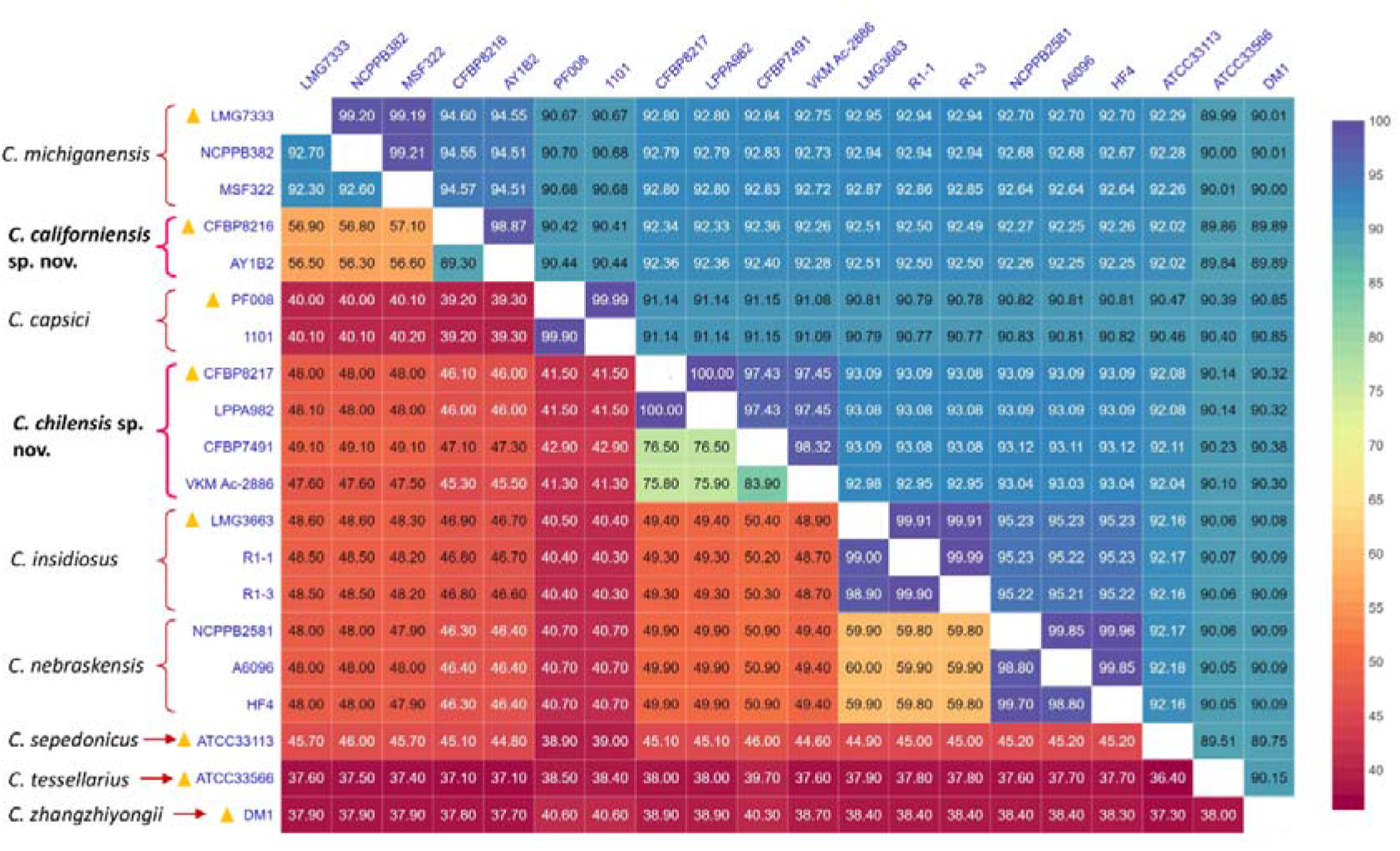
Combined average nucleotide identity (ANI, upper diagonal) and digital DNA-DNA hybridization (dDDH, lower diagonal) heatmap among all *Clavibacter* species. Cut-off values for species delineation are 96% (ANI) and 70% (dDDH).

### Genome Features

To provide a correct taxonomic delineation of the remaining *Clavibacter* subspecies, the complete genomes of type strains of *C. michiganensis* subsp. *californiensis*, *C. michiganensis* subsp. *chilensis*, *C. michiganensis* subsp. *phaseoli*, along with the type strain of *C. tessellarius* and *C. nebraskensis* strain A6096 were sequenced using PacBio RS II (Pacific Biosciences Inc., Menlo Park, CA, United States) powered by single-molecule real-time (SMRT) technology. Genome sequencing was carried out at the Washington State University facility. For whole genomic DNA extraction, a single colony corresponding to each *Clavibacter* strain was streaked on YSC medium [11] and grown overnight at 28°C. Later, half loopful of pure bacterial culture was picked and genomic DNA was extracted using the QIAGEN Genomictip 100/G (Qiagen, Valencia, CA, United States) following the manufacturer’s instructions. DNA was re-suspended into 300-350 μL TE (Tris-EDTA) buffer pH 8.0. Quantification and quality of DNA was analyzed by using NanoDrop spectrophotometer, Qubit 4, and by electrophoresis on 1.5 % agarose gels. Genomic DNA libraries were prepared with a 20 kb insert size and sequenced using P6 polymerase enzyme and C4 sequencing chemistry. Prior assembly, long sequencing reads were trimmed according to the length and quality to obtain highly accurate long reads. Final assembly was done using the Hierarchical Genome Assembly Process (HGAP) version 4.0 (Pacific Biosciences, SMRT Analysis Software v2.3.0). The generated high-quality complete genomes of the five *Clavibacter* strains were annotated in three different pipelines: the NCBI Prokaryotic Genome Annotation Pipeline (PGAP) version 4.8 [26] the Integrated Microbial Genomes annotation pipeline version v.5.0.3 [27, 28] from the Joint Genome Institute (IMG-JGI) and the Rapid Annotation System Technology (RAST) server [29]. All genomes were deposited in the NCBI GenBank genome database with the accession numbers: CP040792-CP040794 (*C. michiganensis* subsp. *californiensis* CFBP 8216ᵀ) CP040795-CP040796 (*C. michiganensis* subsp. *chilensis* CFBP 8217ᵀ), CP040786-CP040787 (*C. michiganensis* subsp. *phaseoli* LPPA 982ᵀ), CP040788-CP040791 (*C. tessellarius* ATCC 33566ᵀ) and CP040797 (*C. nebraskensis* A6096).

The genomes size of *C. michiganensis* subsp. *californiensis* CFBP 8216ᵀ, *C. michiganensis* subsp. *chilensis* CFBP 8217ᵀ, *C. michiganensis* subsp. *phaseoli* LPPA 982ᵀ, *C. tessellarius* ATCC 33566ᵀ and *C. nebraskensis* A6096 consisted of 3,255,537 bp with 3,032 coding sequences (CDS), 3,221,227 bp with 3,016 CDS, 3,22,6754 with 3,021 CDS, 3,366,694 with 3,145 CDS and 3,069,018 with 2,884 CDS, respectively. All five genomes contained six rRNA genes (2 copies of 5S rRNA, 2 copies of 16S rRNA and 2 copies of 23S rRNA), 45 tRNA genes and three non-coding RNA genes. The DNA G+C contents of the genomes were 72.78%, 73.43%, 73.42%, 73.63%, and 72.97%, respectively. Strains CFBP 8216ᵀ harbored 2 plasmids of 78,006- and 20,294-bp while strains CFBP 8217ᵀ and LPPA 982ᵀ presented a single plasmid of 117,510- and 123061-bp, respectively. ATCC 33566ᵀ harbored three plasmids with lengths of 45,791-, 37,356- and 29,539-bp. No plasmid was found in *C. nebraskensis* A6096. Detail description of different genome features for each of the five bacterial strains sequenced in this study are provided in Table 1.

**Table 1.** Genome features and assembly statistics for the five *Clavibacter* strains sequenced in this study.

| Genomic Features | *C. michiganensis* subsp. *californiensis* CFBP 8216<sup>T</sup> | *C. michiganensis* subsp. *chilensis* CFBP 8217<sup>T</sup> | *C. michiganensis* subsp. *phaseoli* LPPA 982<sup>T</sup> | *C. tessellarius* ATCC 33566<sup>T</sup> | *C. nebraskensis* A6096 |
| --- | --- | --- | --- | --- | --- |
| GenBank accession no. | CP040792-CP040794 | CP040795-CP040796 | CP040786-CP040787 | CP040788-CP040791 | CP040797 |
| GOLD project ID in IMG database | Gp0434066 | Gp0434065 | Gp0432876 | Gp0432876 | Gp0434073 |
| Assembly level | Complete | Complete | Complete | Complete | Complete |
| Genome coverage (x) | 712.0x | 373.0x | 468.0x | 574.0x | 289.0x |
| Genome size (bp) | 3243492 | 3221227 | 3226754 | 3329338 | 3069018 |
| G+C content (mol%) <sup>φ</sup> | 72.78 % | 73.43 % | 73.42 % | 73.63 % | 72.97 % |
| Chromosome length (bp) | 3165486 | 3103717 | 3103693 | 3254008 | 3069018 |
| Plasmid length (bp)* | 78006 (pCCa) | 117510 (pCh) | 123061 (pCP) | 29539 (pCT1)<br>45791 (pCT2) | - |
| Genes (total) | 3086 | 3070 | 3075 | 3199 | 2938 |
| Coding sequences (CDS) | 3032 | 3016 | 3021 | 3145 | 2884 |
| Protein coding genes with function prediction <sup>φ</sup> | 2337 | 2397 | 2402 | 2375 | 2229 |
| Protein coding genes with enzymes <sup>φ</sup> | 871 | 870 | 874 | 883 | 858 |
| Protein coding genes connected to KEGG pathways <sup>φ</sup> | 935 | 930 | 932 | 937 | 901 |
| Protein coding genes with COGs <sup>φ</sup> | 2352 | 2420 | 2425 | 2401 | 2256 |
| rRNAs (5S, 16S, 23S) | 2, 2, 2 | 2, 2, 2 | 2, 2, 2 | 2, 2, 2 | 2, 2, 2 |
| tRNAs | 45 | 45 | 45 | 45 | 45 |
| ncRNAs | 3 | 3 | 3 | 3 | 3 |
| Pseudogenes | 67 | 32 | 29 | 100 | 83 |

<sup>φ</sup> Genomic features as annotated using the Integrated Microbial Genomes annotation pipeline v.5.0.3 from the Joint Genome Institute (IMG-JGI).
\*Assigned name for the plasmids is indicated between parentheses.
GOLD: Genomes OnLine Database; KEGG: Kyoto Encyclopedia of Genes and Genomes; COG: Cluster of Orthologous Groups of proteins; rRNA: ribosomal RNA; tRNA: transfer RNA; ncRNA: non-coding RNA.

The average nucleotide identity (ANI) measures the genetic relatedness between a given set of genomes, and it is an important whole genome-based *in silico* tool applied extensively to define species boundaries among prokaryotic genomes [18, 19, 30]. Conversely, the digital DNA-DNA hybridization (dDDH) represents the genome-to-genome distances between complete or incomplete genomic sequences, and it has emerged as a potential bioinformatic method that emulates the traditional wet-lab DNA-DNA hybridization (DDH) employed in prokaryotes taxonomy [21, 31]. In this study, pairwise genome comparisons using ANI tool were computed using the CLC Genomics Workbench 21.0.5 (Qiagen, Germantown, MD) whereas the dDDH was measured using the genome-to-genome distance calculator (GGDC) 3.0 server (http://ggdc.dsmz.de/ggdc.php#, [32]) with the recommended settings: BLAST+ as local alignment tool and formula 2, applied when incomplete genomes are submitted to the server and which consists on the sum of identities within high-scoring segment pairs (HSPs) per total HSP length [21]. ANI and dDDH values were combined in a single matrix and visualized as a heatmap (Figure 1) using the web-tool DISPLAYR (https://www.displayr.com/). The analysis included the five genomes sequenced in this study, along with 15 genomes of the other *Clavibacter* species with their respective type strains. To assign a correct taxonomy status for the remaining *Clavibacter* subspecies along with the other strains examined in this article, the species-delineation framework was set to 96% for ANI [18, 19] and 70 % for dDDH [33]. Tables S2 and S3 show the ANI and dDDH values calculated among 79 *Clavibacter* strains. The strain CFBP 2404ᵀ was not included in the analyses since this isolate is the same type strain LMG 3663ᵀ.

Overall, pairwise ANI and dDDH values among all *Clavibacter* species were lower than the recommended species threshold of 96% [18, 19] and 70% [33], respectively, supporting the elevation of some subspecies to the species level [12]. When comparing with other *Clavibacter* species, *C. tessellarius* and *C. zhangzhiyongii* were the most divergent species within the *Clavibacter* genus with the lowest ANI value of 90% for both, and dDDH of 38% and 40%, respectively. Interspecies ANI numbers were observed less than 91% and 90% for pepper and potato pathogens, *C. capsici* and *C. sepedonicus*, respectively, whereas the minimum dDDH calculations were 38% and 36%, respectively. In agreement with the MLSA phylogenetic tree, *C. nebraskensis* and *C. insidiosus* showed to be closely related since a high ANI of 95% and a dDDH of 60% were obtained between these two species. Moreover, the minimal interspecies ANI/dDDH values for *C. sepedonicus*, *C. insidiosus* and *C. nebraskensis* were 92-/46-, 91-/40- and 91-/41-%, respectively. The highest ANI and dDDH values between *C. michiganensis* and the other *Clavibacter* species or subspecies were 94.5% and 56.3%, respectively, below the proposed 96% ANI and 70% dDDH for the bacterial species boundary, corroborating the recent emended nomenclature of this species [13]. *Clavibacter michiganensis* subps. *californiensis* CFBP 8216ᵀ exhibited ANI and dDDH values of 98.87% and 89.3%, respectively, with respect to the strain AY1B2, isolated from ryegrass, indicating that these two strains belong to the same species. On the other hand, the ANI values of CFBP 8216ᵀ and AY1B2 with the other species within the *Clavibacter* genus varied from 94.60% to 89.84%, while dDDH values ranged 57.10- to 37.10-%, which were below the cut-off parameters for species delineation, 96% ANI and 70% dDDH, revealed that *C. michiganensis* subsp. *californiensis* must be elevated at the species level, and also, the taxonomic name of strain AYB2, wrongly described as *C. michiganensis* in the NCIBI database, must be corrected to *C. californiensis* sp. nov. Intriguingly, the type strains of *C. michiganensis* subsp. *chilensis* CFBP 8217ᵀ and *C. michiganensis* subsp. *phaseoli* LPPA 982ᵀ shared the highest nucleotide similarity of 100% for both ANI and dDDH, indicating that these two bacteria must be merged as a single species, as suggested in previous publications [23, 24]. In addition, ANI and dDDH analyses of strains CFBP 7491, VKM Ac-2886 with CFBP 8217ᵀ and LPPA 982ᵀ revealed average values above the cut-off for species delineation (ANI 98%; dDDH 78%), indicating that all four strains belong to the same species. Interspecies ANI and dDDH values between these four isolates and all other *Clavibacter* species ranged 93-90% and 50-38%, respectively, which are below the edge of species threshold, denoting that these strains represent a different species within the genus *Clavibacter*. Hence, we proposed the name of *C. chilensis* sp. nov. with CFBP 8217ᵀ as the type strain, as formerly published by Yasuhara-Bell and Alvarez [11]. Although *C. michiganensis* subsp. *phaseoli* LPPA 982ᵀ was reported as the causal agent of bean leaf yellowing [9], no symptoms were induced in recent pathogenicity assays performed by Osdagui *et al*. [23] using this pathogen and *C. michiganensis* subsp. *chilensis* CFBP 8217ᵀ on common bean (cv. Red Kideny, cv. Pinto and cv. Navy), cowpea, mung bean, tomato, and pepper. Since both CFBP 8217ᵀ and LPPA 982ᵀ were demonstrated to be the same species [23, 24, this study] and CFBP 8217ᵀ was named based on its geographic origin of isolation instead of the host like LPPA 982ᵀ, we consider that *C. chilensis* sp. nov. is the best appropriate name for describing the four above discussed strains, which have been isolated from distinct hosts such as tomato seeds (CFBP 8217ᵀ and CFBP 7491), bean seeds (LPPA 982ᵀ) and red elderberry (VKM Ac-2886).

Non-pathogenic and seed-associated *Clavibacter*, tomato strains, have shown a wide genetic diversity, and phylogenetically different from the pathogenic *C. michiganensis* strains [11, 25, 34–36]. Moreover, several strains isolated from surface-disinfected tomato parts have been frequently described as endophytic *Clavibacter* strains with no disease symptoms, but shown colonization inside the tomato vascular tissues [2, 25]. Thapa *et al*. [25] reported five endophytic tomato *Clavibacter* strains (CASJ009, CFBP 7494, CFBP 7576, CFBP 8017, and CFBP 8019)—based on phylogenomic analysis, the strains displayed high heterogeneity and distributed in different clades; and particularly, CFBP 7494, CFBP 7576 and CFBP 8017 clustered with the alfalfa, pepper and wheat *Clavibacter* strains, respectively. Stem canker and mosaic disease symptoms were observed on pepper and wheat plants when inoculated with strains CFBP 7576 and CFBP 8017, respectively [25]. Here, we have assessed the taxonomy position of these three isolates (Tables S2 and S3). Intraspecies ANI (98%) and dDDH (80%) analysis demonstrated that CFBP 7576 belongs to *C. capsici*. Conversely, strains CFBP 7494 and CFBP 8017 exhibited ANI values of 96% (on the edge of species delineation) and 94.9%, respectively, whereas the dDDH values (64.6% and 57.7%, respectively) were lower than the limit for species delineation after comparing with the type strains of *C. insidiosus* and C. *tessellarius*, respectively,—demostrated that both bacteria are closely related to the alfalfa and wheat strains, but genetically constitute novel species. This is in sync with our MLSA results, where both isolates formed a separate like-outgroup clade within *C. insidiosus* and *C. tessellarius* clusters (Figure 2). Interestingly, the ANI value between strain DOAB 609, described as *C. tessellarius* in the NCBI database, and *C. tessellarius* ATCC 33566ᵀ was 93.17%, while dDDH value was 49%; indicating that the taxonomic name of this strain is incorrectly assigned. Moreover, DOAB 609 showed to be closely related to the tomato endophyte CFBP 8017 based on our phylogenetic analysis (Figure 2); however, the ANI and dDDH data (92.78% and 47.2%, respectively) between these strains revealed they are different species. Additionally, ANI and dDDH values between DOAB 609 and other *Clavibacter* members ranged 89-90% and 36.2-39.5%, respectively, highlighted that this strain is a novel *Clavibacter* species. On the other hand, ANI and dDDH scores well above the species threshold (98.57% and 85.4%, respectively) were calculated between CFBP 3399, isolated from wild tulip, and the type strain of *C. tessellarius*, corroborating that the tulip strain belongs to the species *C. tessellarius*, which is coherent with the obtained MLSA phylogeny (Figure 2).

**Figure 2.**
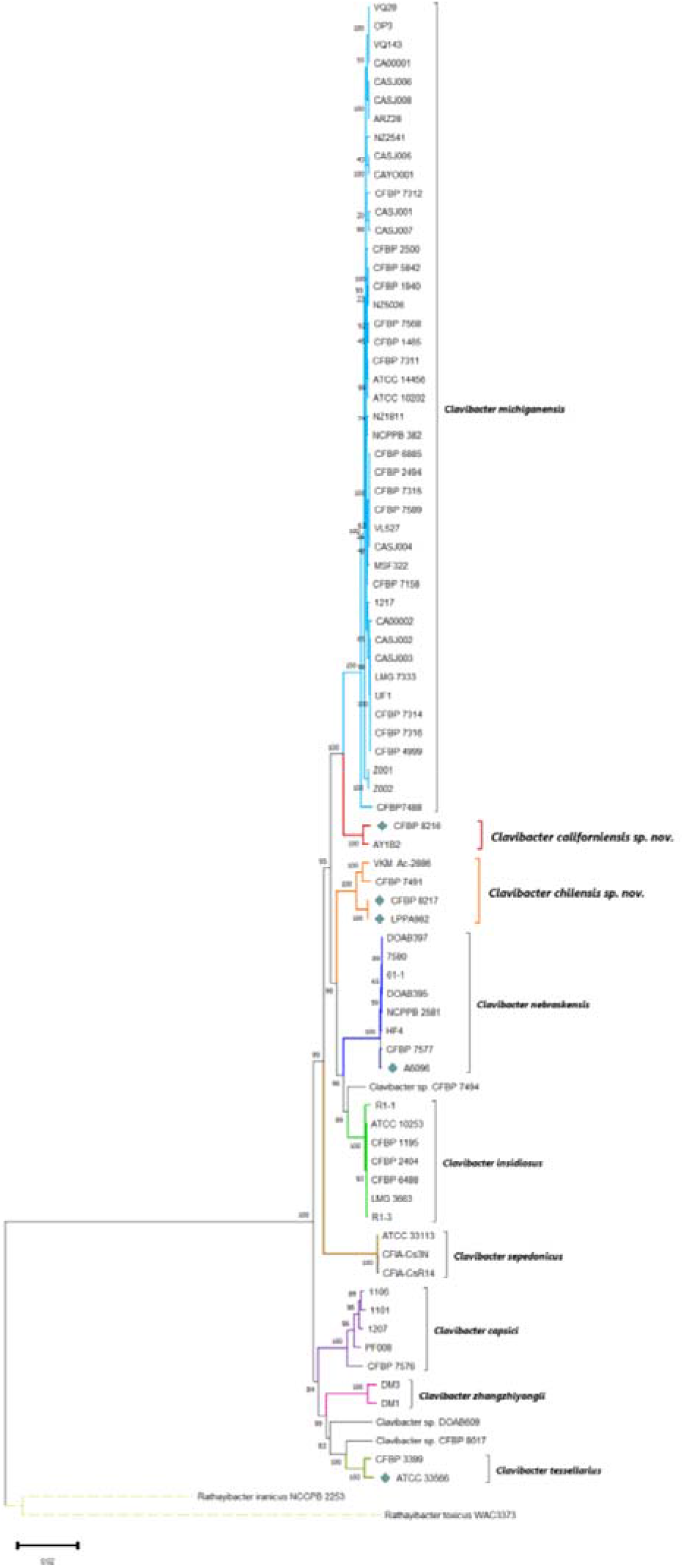
Multi-Locus Sequence Analysis (MLSA) based on the concatenated alignment of seven housekeeping genes, *acnA*, *atpD*, *bipA*, *icdA*, *mtlD*, *recA*, and *rpoB*. The maximum-likelihood (ML) phylogenetic tree was created using 80 *Clavibacter* strains with a bootstrap test of 1,000 replicates. Bootstrap values are indicated at branch points. Evolutionary distances were estimated by applying the Maximum Composite Likelihood (MCL) approach. There was a total of 13,154 positions (alignment length) in the final dataset. The tree was drawn to scale, with branch lengths measured in the number of substitutions per site. The branches were color-coded to emphasize the clusters of each species. The genomes of the five *Clavibacter* strains sequenced in this study are pointed out with a teal diamond. *Rathayibacter iranicus* NCPPB 2253 and *R. toxicus* WAC3373 served as outgroups to root the tree. The evolutionary analysis was conducted in MEGA 11.

To further examine the taxonomy position of a large panel of *Clavibacter* strains including the five genomes sequenced here, a phylogenetic tree based on 1,028 core genes was generated (Figure 3). The FASTA genome files of 78 *Clavibacter* strains were re-annotated using the Rapid prokaryotic genome annotation pipeline Prokka [37]. The genome of *C. nebraskensis* CFBP 7577 was excluded due to the excessive number of contigs and to avoid interferences during the identification of core genes. The annotated GFF3 assemblies obtained using Prokka were applied to perform a pan-genome analysis using the ROARY pipeline [38]. A multiFASTA alignment based on all core genes derived from the pan-genome was created using PRANK [39]. The core-genome alignment served to construct a maximum likelihood (ML) phylogenetic tree using RAxML-NG [40] with a General Time-Reversible GAMMA model and bootstrap test of 1,000 replicates. Lastly, the phylogenetic ML tree was plotted, mid-point rooted, arranged according to increase order of nodes and color-coded using FigTree v1.4.4 (http://tree.bio.ed.ac.uk/software/figtree/).

**Figure 3.**
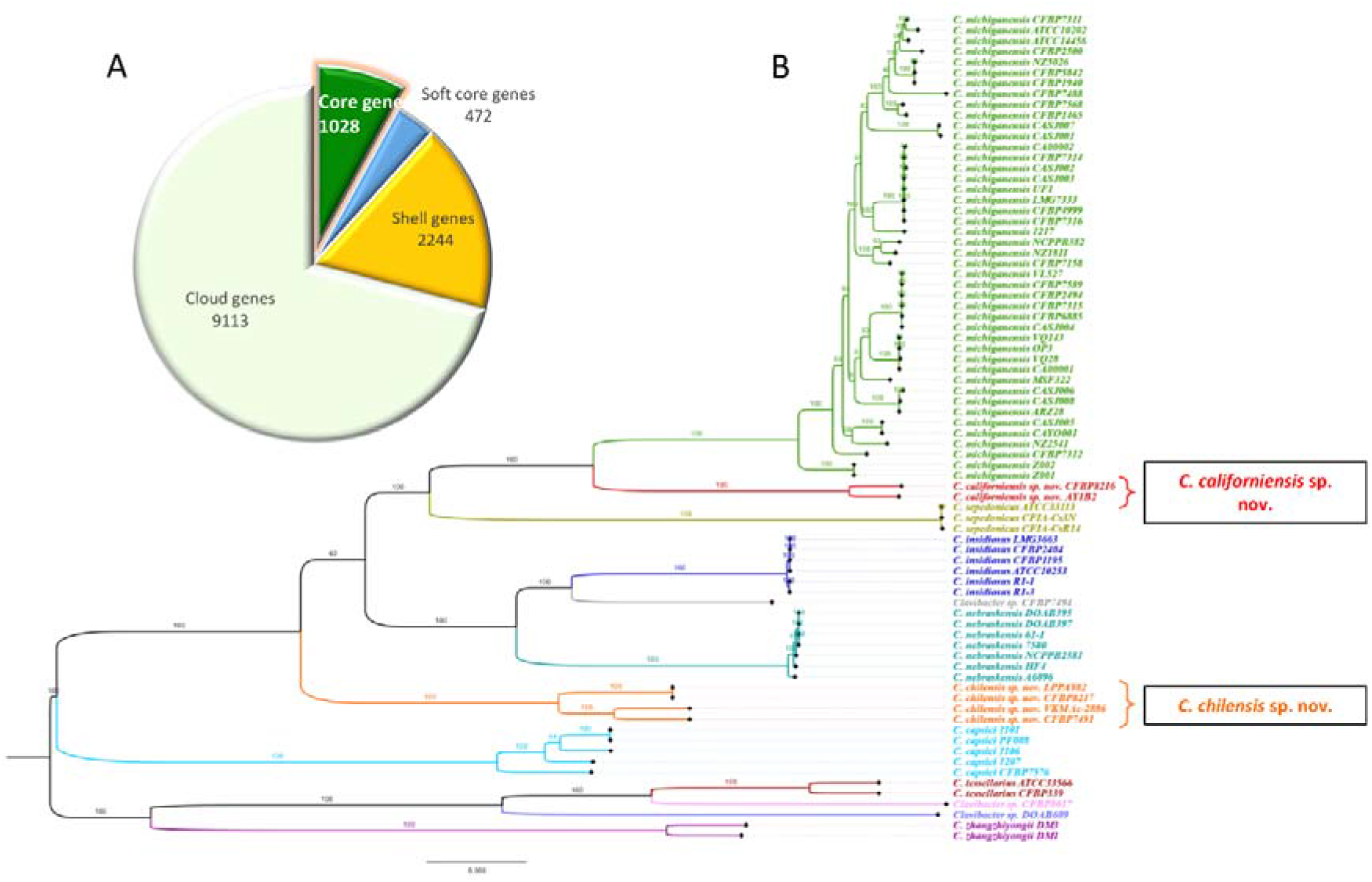
Maximum-likelihood (ML) phylogeny generated from the core-genome of 78 *Clavibacter* strains. (A) Pie chart depicting the proportion of core and accessory genes among the 78 *Clavibacter* genomes as predicted by the ROARY pipeline. (B) ML phylogentic tree reconstructed from the concatenated alignment of 1,028 core genes. A bootstrap test of 1,000 replicates was used to assess the statistical support of each node. The ML analysis was performed using the RAxML-NG tool with a General Time-Reversible GAMMA model. The phylogenetic tree was mid-point rooted and illustrated based on the increasing order of nodes. All strains and their respective branches of each *Clavibacter* species are highlighted with specific colors. The proposed species are pinpointed in a black rectangular frame. FigTree v1.4.4 was used to visualize and color-coded the ML tree.

In sync with the MLSA analysis, the core-genome phylogeny showed 9 clusters that represented each of the *Clavibacter* species. The cluster of *C. californiensis* sp. nov. included the type strain CFBP 8216ᵀ and the ryegrass strain AY1B2. Even though *C. californiensis* sp. nov. positioned closely to the *C. michiganesis* group, *C. californiensis* formed its own distinct and exclusive cluster, thus supporting their elevation to species level. The type strains of *C. michiganensis* subsp. *chilensis* CFBP 8217ᵀ and *C. michiganensis* subsp. *phaseoli* LPPA 982ᵀ clustered far away from the tomato pathogen strains—the branches of both strains converged from the same node, corroborating our previous analysis that this both subspecies must be elevated to the species level and merged as single species, for which we proposed the name *C. chilensis* sp. nov [11]. Additionally, the seed-associated tomato strain CFBP 7491 and the strain VKM Ac-2886 grouped together with CFBP 8217ᵀ and LPPA 982ᵀ, revealing that *C. chilensis* sp. nov. harbors both pathogenic and endophytic strains isolated from different hosts. Importantly, the reconstructed core-genome ML tree confirmed that the tulip strain CFBP 3399 belongs to *C. tessellarius* since the nodes of this strain, along with ATCC 33566ᵀ clustered together. Moreover, an outgroup-like pattern was observed for the nodes of strains CFBP 7494 and CFBP 8017, which positioned close to *C. insidiosus* and *C. tessellarius*, showing that these strains are different species but are related to the alfalfa and wheat strains, respectively. In concordant with the MLSA, ANI and dDDH data, strain DOAB 609 clustered in its own clade and a bit distant from *C. tessellarius* ATCC 33566ᵀ and CFBP 8017, corroborating that DOAB 609 is misnamed as *C. tessellarius*; and hence, a correct name must be given to this new species. An identical clustering pattern of well-defined clades containing each of the *Clavibacter* species was visualized in the ANI-based phylogeny (Figure 4) derived from the computed ANI (Table S2), emphasizing the proposal of *C. californiensis* sp. nov. and *C. chilensis* sp. nov. as species. The ANI phylogenetic tree was built from the whole-genome alignment of 79 *Clavibacter* strains using the neighbor joining method with a bootstrap test of 1,000 replicates (Figure 4). The genome of *R. toxicus* WAC3373 was used as an outgroup and the phylogenetic analysis was generated in the CLC Genomics Workbench v.21.0.5.

**Figure 4.**
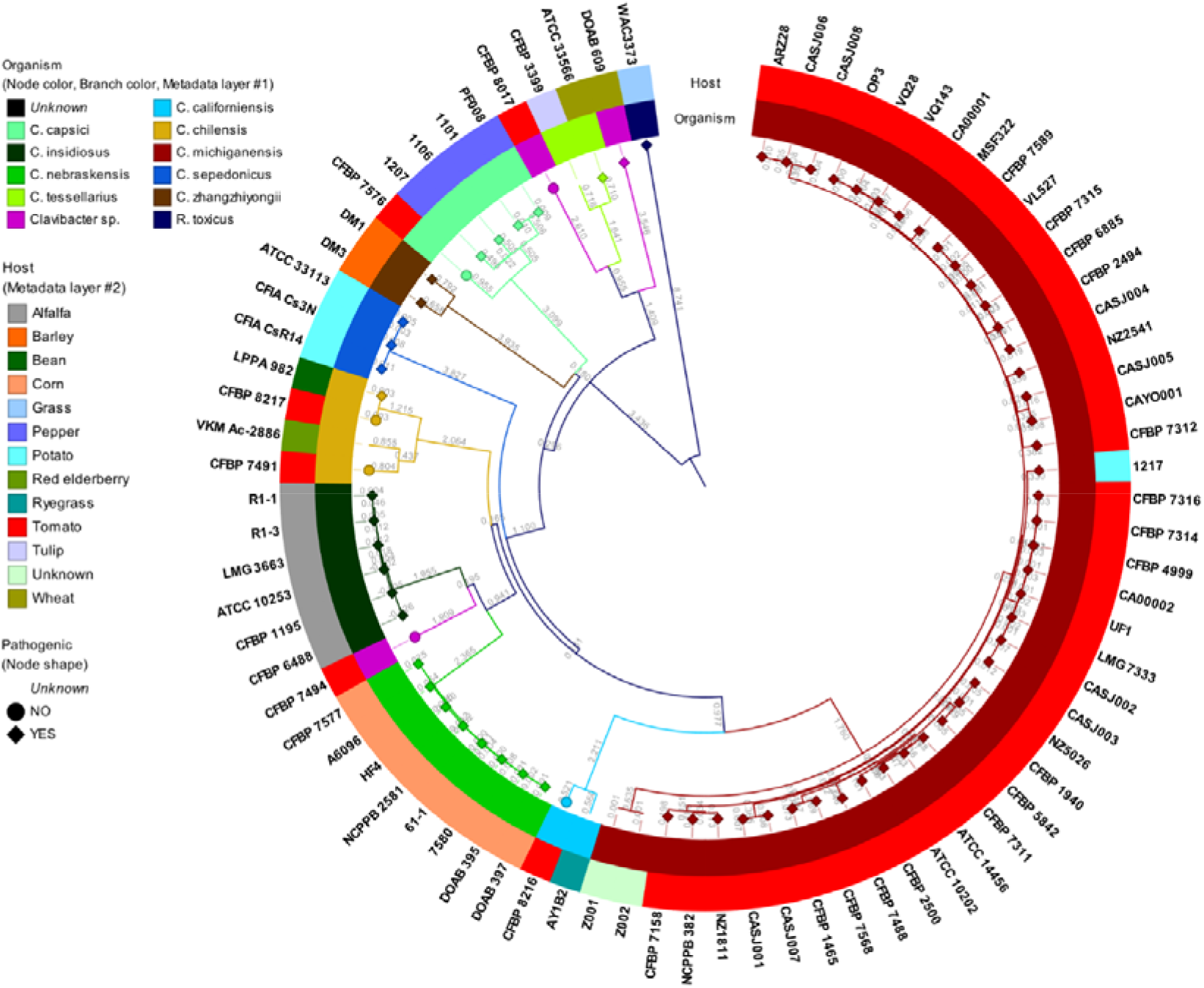
Whole-genome phylogeny based on the computed average nucleotide identity (ANI) of 79 *Clavibacter* strains. The neighbor joining tree was created using the CLC Genomics Workbench v.21.0.5 with a bootstrap test of 100 replicates. *Rathayibacter toxicus* WAC3373 was used as an outgroup to root the tree. The legends of layers corresponding to the organism and host, along with the node shape, are indicated on the left side of the graphic.

The type strain genome server (TYGS; https://tygs.dsmz.de/) was used to confirm the taxonomy status of strains CFBP 7576, CFBP 7491, VKM Ac-2886, CFBP 7494, CFBP 8017 and DOAB 609. TYGS is a high-throughput platform that implements the techniques of the GGDC and compares the submitted genomes with the most curated, updated and valid database of type strain genomes [32, 41]. The web server not only performs a phylogenomic analysis based on the computed dDDH, but also it allows to delineate species according to the established 70% dDDH [32, 41]; and hence, this is a reliable approach for microbial classification. In correlation with our previous analysis, TYGS categorized the strains CFBP 7494, CFBP 8017 and DOAB 609 as potential new species while strain CFBP 7576 was presented as *C. capcisi* and strains CFBP 7491 and VKM Ac-2886 were identified as *C. michiganensis* subsp. *chilensis* (here proposed as *C. chilensis* sp. nov.). A more elaborated taxonomy study is required to provide the most idoneous distinctive name for the strains CFBP 7494, CFBP 8017 and DOA B609, inferred here as novel *Clavibacter* species.

### Orthologous Analysis

Since *C. michiganensis* subsp. *chilensis* CFBP 8217ᵀ and *C. michiganensis* subsp. *phaseoli* LPPA 982ᵀ shared the highest nucleotide identity in the ANI and dDDH (both 100%) pairwise analyses, we performed a dedicated comparative orthologous assessment using the OrthoMCL pipeline v1.4 [42] between these two strains. Homologous protein sequences were computed based on an all-against-all BLASTp search approach with an E-value threshold <1 x 10^−05^, 90% similarity and an alignment coverage >70 %. The precalculated sequence similarity matrix was subjected to a Markov Cluster (MCL) algorithm analysis to determine orthologs groups using a default inflation value of 1.5. A total of 2,873 and 2,874 coding genes were predicted by OrthoMCL in the chromosomic genomes of CFBP 8217ᵀ and LPPA 982ᵀ, respectively. The number of core genes identified between the chromosomes of both strains was 2,857 while only one unique gene (FGI33_12335) encoding for a cysteine hydrolase was found in LPPA 982ᵀ (Figure 5A). In the orthologous analysis between the plasmids of both strains, 114 and 110 genes were found in LPPA 982ᵀ and CFBP 8217ᵀ, respectively. The core gene sets contained between the plasmids were 107 whereas 3 CDS (FGI33_14985, FGI33_14995 and FGI33_15000 encoding for a helix-turn-helix transcriptional regulator, arsenate reductase ArsC and flavoprotein, respectively) were identified as unique genes in the plasmid of LPPA 982ᵀ (Figure 5A). The high number of orthologous genes observed between the genomes of strains CFBP 8217ᵀ and LPPA 982ᵀ corroborates not only that both strains belong to the same species but also that these two isolates seem to be clonal. All defined orthologous genes in the chromosome and plasmid of CFBP 8217ᵀ and LPPA982ᵀ were illustrated in a Circos plot (Figure 5B) [43].

**Figure 5.**
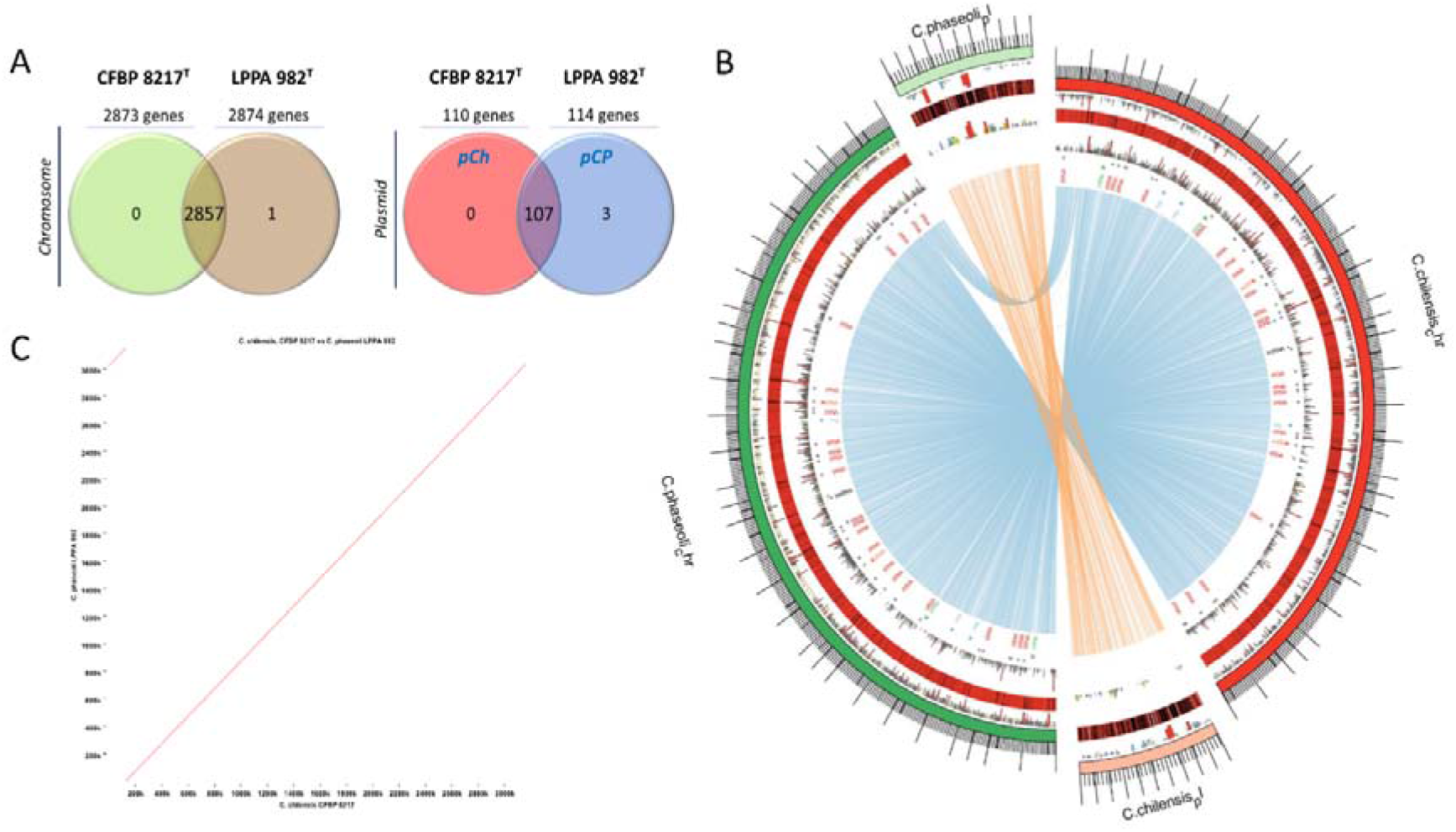
Orthologous analysis between *C. michiganensis* subsp. *chilensis* CFBP 8217ᵀ and *C. michiganensis* subsp. *phaseoli* LPPA 982ᵀ. (A) Venn diagrams based on the core-genome generated from the OrthoMCL analysis between the chromosome and plasmid of CFBP 8217ᵀ and LPPA 982ᵀ. The top panel over each circle indicates the total number of genes found in each strain at the chromosome and plasmid levels. The value located in the intersection of both circles shows the number of shared genes (core genes) between the chromosomes and plasmids of both strains. (B) Circos plot [43] illustrating the orthologous genes found between the chromosomes and plasmids of both strains. From the outermost to the innermost the layers portray: the genome coordinates (axis in kb) of the chromosomes and plasmids; bar chart illustrating the sizes of those loci with a negative orientation; a red ring showing with black lines all coding sequences found in the chromosome and plasmid of each strain (the genes of the highest length can be visualized with a clear black line); bar chart illustrating the sizes of those loci with a positive orientation; text description pointing out the location of tRNA (red), rRNA (green), mRNA (black), ncRNA (orange) and regulatory genes (blue) in the chromosomes of both strains; and the inner layer depicting the connections among all homologous genes predicted between the chromosomes and plasmids of CFBP 8217ᵀ and LPPA 982ᵀ. (C) *in-silico* syntenic dot plot showing a high similarity between the genomes of CFBP 8217 ᵀ (x-axis) vs LPPA 982ᵀ (y-axis).

To further evaluate the genome similarities between the type strains of *C. michiganensis* subsp. *chilensis* and *C. michiganensis* subsp. *phaseoli*, an *in-silico* dot plot analysis was performed using the “Create Whole Genome Dot Plot” tool as implemented in the CLC Genomics Workbench v.21.0.5. The syntenic dot plot graphic (Figure 5C) displayed a clear collinear diagonal line between the sequences of strains CFBP 8217ᵀ and LPPA 982ᵀ, indicating that the genomes of both strains are highly identical. This result emphasizes that strains CFBP 8217ᵀ and LPPA 982ᵀ must be considered as a single species.

## Conclusion

To summarize, in the present study, we have sequenced the complete genomes of the type strains of the still-assigned subspecies *C. michiganensis* subsp. *californiensis* CFBP 8216ᵀ, *C. michiganensis* subsp. *chilensis* CFBP 8217ᵀ and *C. michiganensis* subsp. *phaseoli* LPPA982ᵀ along with C. *tessellarius* ATCC 33566ᵀ and *C. nebraskensis* A6096. This manuscript provides conclusive evidence to elevate the remaining three subspecies within the *Clavibacter* genus to the species level. By using different state-of-the-art *in silico* strategies such as MLSA, ANI, dDDH, core-genome phylogenomics and whole-genome based ANI phylogeny, we have demonstrated that *C. michiganensis* subsp. *californiensis* must be raised to a species level for which we propose the name *Clavibacter californiensis* sp. nov. (type strain CFBP 8216ᵀ=ATCC BAA-2691ᵀ =C55ᵀ). Likewise, different comparative genomic methods like orthologous and genome synteny analyses, as well as all above mentioned bioinformatic approaches, pointed out that type strains of *C. michiganensis* subsp. *chilensis* CFBP 8217ᵀ and *C*. *michiganensis* subsp. *phaseoli* LPPA982ᵀ must be merged as a single organism and raised to a species level; consequently, we propose the name *C. chilensis* sp. nov. (type strain CFBP 8217ᵀ = ATCC BAA-2690ᵀ = ZUM3936ᵀ). Additionally, we clarified the taxonomy description of a large panel composed of 80 *Clavibacter* strains. Our polyphasic analyses allowed us to determine that the ryegrass isolate AY1B2 is another strain of the proposed species *C. californiensis* sp. nov. while strains CFBP 7491 and VKM Ac-2886 isolated from tomato seeds and red elderberry, respectively, must be re-classified as *C. chilensis* sp. nov. Furthermore, we corroborated that the tomato endophyte strain CFBP 7576 constitutes another member of *C. capsici*. Interestingly, our high-throughput *in-silico* assessments revealed that the tulip strain CFBP 3399 belongs to the wheat pathogenic species, *C. tessellarius*. On the other hand, analysis indicated that strain DOAB 609 has been misnamed as *C. tessellarius* in the NCBI GenBank database, and should be described as a novel species of *Clavibacter*. In a similar way, the tomato endophytic strains CFBP 7494 and CFBP 8017 must be considered as a new species according to the calculated ANI and dDDH scores. Lastly, the TYGS allowed us to conclude that strains DOAB 609, CFBP 7494 and CFBP 8017 are certainly belong to new species. A further elaborated taxonomy study must be undertaken to provide the best fitting name for these potential novel species.

## DESCRIPTION OF *CLAVIBACTER CALIFORNIENSIS* SP. NOV

Elevation in taxonomic rank from *Clavibacter michiganensis* subsp. *californiensis* (Yasuhara-Bell and Alvarez, 2015) to *Clavibacter californiensis* (ca.li.for.ni.en′sis. N.L. masc. adj. *californiensis* pertaining to California, referring to the location of isolation of the type strain).

Basonym: *Clavibacter michiganensis* subsp. *californiensis* Yasuhara-Bell and Alvarez 2015.

The description of this taxon remains the same from its first description as *Clavibacter michiganensis* subsp. *californiensis* reported by Yasuhara-Bell and Alvarez, 2015.

The type strain is C55ᵀ (=CFBP 8216ᵀ=ATCC BAA-2691ᵀ) and it was isolated from seeds of *Solanum lycopersycum* in the state of California, USA. The DNA G+C content of the type strain was 72.78% -while the DNA coding number of bases was 91.57%.

## DESCRIPTION OF *CLAVIBACTER CHILENSIS* SP. NOV

Merging of *Clavibacter michiganensis* subsp. *phaseoli* (Gonzalez and Trapiello, 2014) and *Clavibacter michiganensis* subsp. *chilensis* (Yasuhara-Bell and Alvarez, 2015) as a single species and elevate to taxonomic level as *Clavibacter chilensis* (chil.en′sis. N.L. masc. adj. *chilensis* pertaining to Chile, referring to the location of isolation of the type strain).

Basonym: *Clavibacter michiganensis* subsp. *chilensis* Yasuhara-Bell and Alvarez 2015, *Clavibacter michiganensis* subsp. *phaseoli* Gonzalez and Trapiello 2014.

The description of this species is unchanged from its former information as *Clavibacter michiganensis* subsp. *chilensis* provided by Yasuhara-Bell and Alvarez, 2015.

The type strain is ZUM3936ᵀ ( = ATCC BAA-2690ᵀ = CFBP 8217ᵀ). The bacterium was isolated from seeds of *Solanum lycopersycum* from Chile. The DNA G+C content of the type strain is 73.43% whereas the DNA coding number of bases is 92.04%.

## AUTHOR STATEMENTS

### Ethical approval

*N/A*

### Consent for publication

*N/A*

## SUPPLEMENTARY DATA

**Table S1.** Detailed list of genomic information of the 76 *Clavibacter* strains retrieved from the NCBI GenBank database and the five *Clavibacter* genomes sequenced in this study (highlighted in light blue color). The strains presented in this table were used for different *in silico* analyses such as MLSA, ANI and *dDDH* computation, reconstructed ML phylogenomic tree from core-genome alignment, ANI-based phylogeny, and taxonomy assessment of undefined species in the TYGS platform.

**Table S2.** Pairwise heatmap comparison based on the ANI computation values among the 79 *Clavibacter* strains, including the five strains sequenced in this study. The ANI scores were determined in the CLC Genomics Workbench 21.0.5. Strain CFBP 2404ᵀ was not included in the analysis since this isolate is the same type strain LMG 3663ᵀ

**Table S3.** Pairwise heatmap comparison based on the dDDH computation values among the 79 *Clavibacter* strains, including the five strains sequenced in this study. The dDDH scores were calculated in the web server Genome-to-Genome Distance Calculator (GGDC) version 3.0 (http://ggdc.dsmz.de/ggdc.php#). Strain CFBP 2404ᵀ was not included in the analysis since this isolate is the same type strain LMG 3663ᵀ

