## Supplemental Table 1 for "Elevation of *Clavibacter michiganensis* subsp. *californiensis* to species level as *Clavibacter californiensis* sp. nov., merging and re-classification of *Clavibacter michiganensis* subsp. *chilensis* and *Clavibacter michiganensis* subsp. *phaseoli* as *Clavibacter chilensis* sp. nov. based on c"

### SUPPLEMENTARY DATA

**Table S1.** Detailed list of genomic information of the 76 *Clavibacter* strains retrieved from the NCBI GenBank database and the five *Clavibacter* genomes sequenced in this study (highlighted in light blue color). The strains presented in this table were used for different *in silico* analyses such as MLSA, ANI and *dDDH* computation, reconstructed ML phylogenomic tree from core-genome alignment, ANI-based phylogeny, and taxonomy assessment of undefined species in the TYGS platform.

| Former species name in NCBI | Proposed taxonomy | Strain ID | Biotope/host | Geographic origin | Year isolated | GenBank accession number | Assembly level | Genome size (Mb) | Scaffolds | Sequencing technology | Genome coverage | Assembly method |
|---|---|---|---|---|---|---|---|---|---|---|---|---|
| *Clavibacter michiganensis* subsp. *michiganensis* | *C. michiganensis* | CAYO001 | *Solanum lycopersicum* | California, USA | 2001 | MDHL00000000 | Contig | 3.33 | 1 | Illumina HiSeq; PacBio | 100.0x | SPAdes v. 3.6; SMRT analysis v. 2.2.0 |
| *C. m.* subsp. *michiganensis* | *C. michiganensis* | CASJ002 | *Solanum lycopersicum* | California, USA | 1999 | MDHC00000000 | Contig | 3.28 | 1 | Illumina MiSeq; PacBio | 100.0x | SPAdes v. 3.6; SMRT analysis v. 2.2.0 |
| *C. m.* subsp. *michiganensis* | *C. michiganensis* | LMG 7333[T] | *Solanum lycopersicum* | Hungary | 1957 | MZMP00000000 | Contig | 3.39 | 5 | PacBio | 516.0x | Celera Assembler v. 1 |
| *C. m.* subsp. *michiganensis* | *C. michiganensis* | NCPPB382 | *Solanum lycopersicum* | UK | 1956 | NC_009480 | Complete | 3.40 | - | Sanger ABI 3700 | | PHRAP |
| *C. m.* subsp. *michiganensis* | *C. michiganensis* | MSF322 | *Solanum lycopersicum* | Chile | 2005 | NZ_CP047051 | Complete | 3.40 | - | Hybrid assembly (MiniION/MiSeq) | 139x | Unicycler v. 0.4.7 |
| *C. m.* subsp. *michiganensis* | *C. michiganensis* | UF1 | *Solanum lycopersicum* | Florida, USA | 2012 | NZ_CP033724 | Complete | 3.37 | - | PacBio | 396.0x | Canu v. 1.5 |
| *C. m.* subsp. *michiganensis* | *C. michiganensis* | VL527 | *Solanum lycopersicum* | Chile | 2012 | NZ_CP047054 | Complete | 3.40 | - | Hybrid assembly (MiniION/MiSeq) | 152x | Unicycler v. 0.4.7 |
| *C. m.* subsp. *michiganensis* | *C. michiganensis* | VQ143 | *Solanum lycopersicum* (plant tissue) | Chile | 2000 | NZ_CP076352 | Complete | 3.26 | - | Hybrid assembly (MiniION/MiSeq) | 547x | SPAdes v. 3.14.1 |
| *C. m.* subsp. *michiganensis* | *C. michiganensis* | VQ28 | *Solanum lycopersicum* (plant tissue) | Chile | 1996 | NZ_CP076349 | Complete | 3.36 | - | Hybrid assembly (MiniION/MiSeq) | 556x | Unicycler v. 0.4.8 |

| | | | | | | | | | | | | |
|---|---|---|---|---|---|---|---|---|---|---|---|---|
| *C. m.* subsp. *michiganensis* | *C. michiganensis* | CA00001 | *Solanum lycopersicum* (plant tissue) | California, USA | 2000 | MDHK00000000 | Contig | 3.47 | 107 | Illumina HiSeq | 100.0x | SPAdes v. 3.6 |
| *C. m.* subsp. *michiganensis* | *C. michiganensis* | CA00002 | *Solanum lycopersicum* (plant tissue) | California, USA | 2000 | MDHM00000000 | Contig | 3.28 | 1 | Illumina HiSeq; PacBio | 100.0x | SPAdes v. 3.6; SMRT analysis v. 2.2.0 |
| *C. m.* subsp. *michiganensis* | *C. michiganensis* | CASJ001 | *Solanum lycopersicum* (plant tissue) | California, USA | 1999 | MDHB00000000 | Contig | 3.31 | 6 | Illumina MiSeq; PacBio | 90.0x | SPAdes v. 3.6; SMRT analysis v. 2.2.0 |
| *C. m.* subsp. *michiganensis* | *C. michiganensis* | CASJ003 | *Solanum lycopersicum* (plant tissue) | California, USA | 1999 | MDHD00000000 | Contig | 3.25 | 423 | Illumina HiSeq | 100.0x | SPAdes v. 3.6 |
| *C. m.* subsp. *michiganensis* | *C. michiganensis* | CASJ004 | *Solanum lycopersicum* (plant tissue) | California, USA | 1999 | MDHE00000000 | Contig | 3.31 | 469 | Illumina HiSeq | 100.0x | SPAdes v. 3.6 |
| *C. m.* subsp. *michiganensis* | *C. michiganensis* | CASJ005 | *Solanum lycopersicum* (plant tissue) | California, USA | 2001 | MDHF00000000 | Contig | 3.20 | 313 | Illumina MiSeq | 90.0x | SPAdes v. 3.6 |
| *C. m.* subsp. *michiganensis* | *C. michiganensis* | CASJ006 | *Solanum lycopersicum* (plant tissue) | California, USA | 2002 | MDHG00000000 | Contig | 3.37 | 2 | Illumina MiSeq; PacBio | 100.0x | SPAdes v. 3.6; SMRT analysis v. 2.2.0 |
| *C. m.* subsp. *michiganensis* | *C. michiganensis* | CASJ007 | *Solanum lycopersicum* (plant tissue) | California, USA | 2011 | MDHH00000000 | Contig | 3.35 | 10 | Illumina MiSeq; PacBio | 100.0x | SPAdes v. 3.6; SMRT analysis v. 2.2.0 |
| *C. m.* subsp. *michiganensis* | *C. michiganensis* | CASJ008 | *Solanum lycopersicum* (plant tissue) | California, USA | 2002 | MDHI00000000 | Contig | 3.39 | 32 | Illumina HiSeq | 90.0x | SPAdes v. 3.6 |
| *C. m.* subsp. *michiganensis* | *C. michiganensis* | ARZ28 | *Solanum lycopersicum* (plant tissue) | USA | 2000 | QLNE00000000 | Contig | 3.28 | 434 | Illumina HiSeq | 174.0x | SPAdes v. 3.11 |
| *C. m.* subsp. *michiganensis* | *C. michiganensis* | ATCC 10202 | *Solanum lycopersicum* (plant tissue) | ? | 1946 | QLMX00000000 | Contig | 3.28 | 334 | Illumina HiSeq | 192.0x | SPAdes v. 3.11 |
| *C. m.* subsp. *michiganensis* | *C. michiganensis* | ATCC 14456 | *Solanum lycopersicum* (plant tissue) | Italy | 1961 | QLMU00000000 | Contig | 3.30 | 41 | Illumina HiSeq | 250.0x | SPAdes v. 3.11 |
| *C. m.* subsp. *michiganensis* | *C. michiganensis* | CFBP 1465 | *Solanum lycopersicum* (plant tissue) | France | 1975 | QLMV00000000 | Contig | 3.33 | 66 | Illumina HiSeq | 111.0x | SPAdes v. 3.11 |
| *C. m.* subsp. *michiganensis* | *C. michiganensis* | CFBP 1940 | *Solanum lycopersicum* (plant tissue) | Spain | 1978 | QLMW00000000 | Contig | 3.30 | 35 | Illumina HiSeq | 152.0x | SPAdes v. 3.11 |

| | | | | | | | | | | | | |
|---|---|---|---|---|---|---|---|---|---|---|---|---|
| *C. m.* subsp. *michiganensis* | *C. michiganensis* | CFBP 2494 | *Solanum lycopersicum* (plant tissue) | Algeria | 1984 | QLMY00000000 | Contig | 3.32 | 39 | Illumina HiSeq | 120.0x | SPAdes v. 3.11 |
| *C. m.* subsp. *michiganensis* | *C. michiganensis* | CFBP 2500 | *Solanum lycopersicum* (plant tissue) | Algeria | 1984 | QLMZ00000000 | Contig | 3.31 | 332 | Illumina HiSeq | 139.0x | SPAdes v. 3.11 |
| *C. m.* subsp. *michiganensis* | *C. michiganensis* | CFBP 4999 | *Solanum lycopersicum* (plant tissue) | Hungary | 1957 | RDQW00000000 | Contig | 3.37 | 40 | Illumina HiSeq | 150.0x | Velvet v. 1.2.02; SOAPdenovo v. 2.04 |
| *C. m.* subsp. *michiganensis* | *C. michiganensis* | CFBP 5842 | *Solanum lycopersicum* (plant tissue) | Brazil | 1993 | QLNA00000000 | Contig | 3.34 | 127 | Illumina HiSeq | 95.0x | SPAdes v. 3.11 |
| *C. m.* subsp. *michiganensis* | *C. michiganensis* | CFBP 6885 | *Solanum lycopersicum* (plant tissue) | France | 2004 | QLML00000000 | Contig | 3.33 | 25 | Illumina HiSeq | 139.0x | SPAdes v. 3.11 |
| *C. m.* subsp. *michiganensis* | *C. michiganensis* | CFBP 7158 | *Solanum lycopersicum* (plant tissue) | New Zealand | 1968 | QLNB00000000 | Contig | 3.29 | 23 | Illumina HiSeq | 216.0x | SPAdes v. 3.11 |
| *C. m.* subsp. *michiganensis* | *C. michiganensis* | CFBP 7311 | *Solanum lycopersicum* (plant tissue) | Morocco | 1989 | QLNC00000000 | Contig | 3.30 | 46 | Illumina HiSeq | 142.0x | SPAdes v. 3.11 |
| *C. m.* subsp. *michiganensis* | *C. michiganensis* | CFBP 7312 | *Solanum lycopersicum* (plant tissue) | China | 1998 | QLND00000000 | Contig | 3.32 | 43 | Illumina HiSeq | 216.0x | SPAdes v. 3.11 |
| *C. m.* subsp. *michiganensis* | *C. michiganensis* | CFBP 7314 | *Solanum lycopersicum* (plant tissue) | USA | 2002 | QLMM00000000 | Contig | 3.27 | 30 | Illumina HiSeq | 154.0x | SPAdes v. 3.11 |
| *C. m.* subsp. *michiganensis* | *C. michiganensis* | CFBP 7315 | *Solanum lycopersicum* (plant tissue) | USA | 1998 | QLMN00000000 | Contig | 3.31 | 31 | Illumina HiSeq | 168.0x | SPAdes v. 3.11 |
| *C. m.* subsp. *michiganensis* | *C. michiganensis* | CFBP 7316 | *Solanum lycopersicum* (plant tissue) | USA | 1998 | QLMO00000000 | Contig | 3.26 | 29 | Illumina HiSeq | 138.0x | SPAdes v. 3.11 |
| *C. m.* subsp. *michiganensis* | *C. michiganensis* | CFBP 7488 | *Solanum lycopersicum* (plant tissue) | France | 2008 | QLMP00000000 | Contig | 3.30 | 55 | Illumina HiSeq | 127.0x | SPAdes v. 3.11 |
| *C. m.* subsp. *michiganensis* | *C. michiganensis* | CFBP 7568 | *Solanum lycopersicum* (plant tissue) | USA | 2000 | QLMQ00000000 | Contig | 3.35 | 61 | Illumina HiSeq | 140.0x | SPAdes v. 3.11 |
| *C. m.* subsp. *michiganensis* | *C. michiganensis* | CFBP 7589 | *Solanum lycopersicum* (plant tissue) | Belgium | 1998 | QLMR00000000 | Contig | 3.32 | 35 | Illumina HiSeq | 297.0x | SPAdes v. 3.11 |

| | | | | | | | | | | | | |
|---|---|---|---|---|---|---|---|---|---|---|---|---|
| *C. m.* subsp. *michiganensis* | *C. michiganensis* | NZ1811 | *Solanum lycopersicum* (plant tissue) | New Zealand | 1967 | QLMS00000000 | Contig | 3.27 | 333 | Illumina HiSeq | 170.0x | SPAdes v. 3.11 |
| *C. m.* subsp. *michiganensis* | *C. michiganensis* | NZ2541 | *Solanum lycopersicum* (plant tissue) | United Kingdom | 1962 | QODA00000000 | Contig | 3.32 | 228 | Illumina HiSeq | 193.0x | SPAdes v. 3.11 |
| *C. m.* subsp. *michiganensis* | *C. michiganensis* | NZ5026 | *Solanum lycopersicum* (plant tissue) | USA | 1974 | QLMT00000000 | Contig | 3.28 | 263 | Illumina HiSeq | 120.0x | SPAdes v. 3.11 |
| *C. m.* subsp. *michiganensis* | *C. michiganensis* | 1217 | *Solanum tuberosum* | Russia | 2006 | JAATPM000000000 | Scaffold | 3.31 | 227 | 454 | 526x | GS De Novo Assembler v. 1.1 |
| *C. m.* subsp. *michiganensis* | *C. michiganensis* | OP3 | *Solanum lycopersicum* (plant tissue) | Chile | 2015 | WTCS00000000 | Contig | 3.47 | 5 | Hybrid assembly (MinION; MiSeq) | 139x | Unicycler v. 0.4.7 |
| *C. michiganensis* | *C. michiganensis* | Z001 | ? | ? | ? | PSTW00000000 | Scaffold | 3.30 | 39 | Illumina HiSeq | 112.796x | SPAdes v. 3.7.0 |
| *C. michiganensis* | *C. michiganensis* | Z002 | ? | ? | ? | PSTV00000000 | Contig | 3.32 | 44 | Illumina HiSeq | 160.65x | SPAdes v. 3.7.0 |
| *C. m.* subsp. *californiensis* | *C. californiensis* | CFBP 8216$^{T}$ | *Solanum lycopersicum* (seeds) | California, USA | 2000 | CP040792-CP040794 | Complete | 3.26 | - | PacBio RSII | 712.0x | HGAP v. 4 |
| *C. michiganensis* | *C. californiensis* | AY1B2 ¥ | *Lolium perenne* (ryegrass) | Oregon, USA | 2013 | PSTR00000000 | Scaffold | 3.34 | 42 | Illumina HiSeq | 165.811x | SPAdes v. 3.11.1 |
| *C. m.* subsp. *capsici* | *C. capsici* | PF008$^{T}$ | *Capsicum annuum* (plant stem) | South Korea | 1999 | NZ_CP012573 | Complete | 3.24 | - | PacBio | 67.41x | SMRT Analysis Portal v. 2.2.0 |
| *C. m.* subsp. *capsici* | *C. capsici* | 1101 | *Capsicum annuum* (plant stem) | South Korea | 1997 | NZ_CP048049 | Complete | 3.19 | - | PacBio RSII | 125.04x | HGAP v. 2 |
| *C. m.* subsp. *capsici* | *C. capsici* | 1106 | *Capsicum annuum* (plant stem) | South Korea | 1997 | NZ_CP048047 | Complete | 3.19 | - | PacBio RSII | 296.02x | HGAP v. 2 |
| *C. m.* subsp. *capsici* | *C. capsici* | 1207 | *Capsicum annuum* (plant stem) | South Korea | 1997 | CP048045 | Complete | 3.25 | - | PacBio RSII | 224.88x | HGAP v. 2 |
| *C. michiganensis* | *C. capsici* | CFBP 7576 ¥ | *Solanum lycopersicum* (seeds) | ? | 1997 | MDJX00000000 | Scaffold | 3.39 | 374 | Illumina HiSeq | 90.0x | SPAdes v. 3.6 |
| *C. m.* subsp. *phaseoli* | *C. chilensis* | LPPA 982$^{T}$ | *Phaseolis vulgaris* L. (seeds) | Spain | 2009 | CP040786-CP040787 | Complete | 3.23 | - | PacBio RSII | 468.0x | HGAP v. 4 |

| | | | | | | | | | | | | |
|---|---|---|---|---|---|---|---|---|---|---|---|---|
| *C. m.* subsp. *michiganensis* | *C. chilensis* | CFBP 8217$^{T}$ | *Solanum lycopersicum* (seeds) | Chile | 2007 | CP040795-CP040796 | Complete | 3.22 | - | PacBio RSII | 373.0x | HGAP v. 4 |
| *C. m.* subsp. *phaseoli* | *C. chilensis* | VKM Ac-2886 | *Sambucus racemosa* (plant tissue) | Moscow, Russia | 2017 | JADKRP000000000 | Scaffold | 3.36 | 21 | Illumina NovaSeq | 785x | SPAdes v. 3.14.1 |
| *C. michiganensis* | *C. chilensis* | CFBP 7491 ¥ | *Solanum lycopersicum* (seeds) | ? | ? | QWEB00000000 | Contig | 3.29 | 921 | Illumina HiSeq | 6.0x | Velvet v. 1.2.10 ; SOAPdenovo v. 2.04; SOAPGapCloser v. 1.12 |
| *C. m.* subsp. *insidiosus* | *C. insidiosus* | LMG 3663$^{T}$ | *Medicago sativa* | USA | 1955 | MZMO00000000 | Contig | 3.39 | 3 | PacBio | 458.0x | Celera Assembler v. 1 |
| *C. m.* subsp. *insidiosus* | *C. insidiosus* | R1-1 | *Medicago truncatula* (plant stem) | Minnesota, USA | 2009 | NZ_CP011043 | Complete | 3.41 | - | PacBio RSII SMRT P6-C4, Illumina GAIIx | 163.4x | HGAP3 v. 2.2, Pilon v. 1.10 |
| *C. m.* subsp. *insidiosus* | *C. insidiosus* | R1-3 | *Medicago truncatula* (plant stem) | Minnesota, USA | 2009 | NZ_CP021034 | Complete | 3.39 | - | PacBio RSII | 120x | HGAP3 PacBio SMRT portal v. 2.2.0 |
| *C. m.* subsp. *insidiosus* | *C. insidiosus* | ATCC 10253 | *Medicago sativa* | Kansas, USA | 1960 | NZ_CP021038 | Complete | 3.24 | - | PacBio RSII | 117x | HGAP3 SMRT portal v. 2.2.0 |
| *C. m.* subsp. *insidiosus* | *C. insidiosus* | CFBP 1195 | *Medicago sativa* | United Kingdom | 1964 | QWDZ00000000 | Contig | 3.20 | 805 | Illumina HiSeq | 6.0x | Velvet v. 1.2.10; SOAPdenovo v. 2.04; SOAPGapCloser v. 1.12 |
| *C. m.* subsp. *insidiosus* | *C. insidiosus* | CFBP 6488 ϕ | *Medicago sativa* | Czech Republic | 1998 | QWEA00000000 | Contig | 3.23 | 1892 | Illumina HiSeq | 6.0x | Velvet v. 1.2.10; SOAPdenovo v. 2.04; SOAPGapCloser v. 1.12 |
| *C. m.* subsp. *insidiosus* | *C. insidiosus* | CFBP 2404$^{T}$ | *Medicago sativa* | USA | 1955 | RDQV00000000 | Scaffold | 3.29 | 73 | Illumina HiSeq | 200.0x | Velvet v. 1.2.02; SOAPdenovo v. 2.04 |
| *C. michiganensis* | *Clavibacter* sp. | CFBP 7494 ¥ | *Solanum lycopersicum* (seeds) | ? | | MDJW00000000 | Contig | 3.31 | 15 | Illumina HiSeq | 70.0x | SPAdes v. 3.6 |
| *C. m.* subsp. *nebraskensis* | *C. nebraskensis* | NCPPB 2581$^{T}$ | *Zea mays* | Nebraska, USA | 1974 | NC_020891 | Complete | 3.06 | - | | | |

| | | | | | | | | | | | | |
|---|---|---|---|---|---|---|---|---|---|---|---|---|
| *C. m.* subsp. *nebraskensis* | *C. nebraskensis* | HF4 | *Zea mays* | Iowa, USA | 2012 | NZ_CP033721 | Complete | 3.06 | - | PacBio | 90.0x | HGAP v. can v. 1.5 |
| - | *C. nebraskensis* | A6096 | *Zea mays* | Nebraska, USA | 1971 | CP040797 | Complete | 3.07 | - | PacBio RSII | 289.0x | HGAP v.4 |
| *C. m.* subsp. *nebraskensis* | *C. nebraskensis* | 61-1 | *Zea mays* | Iowa, USA | 2006 | NZ_CP033723 | Complete | 3.07 | - | PacBio | 138.0x | Canu v. 1.5 |
| *C. m.* subsp. *nebraskensis* | *C. nebraskensis* | 7580 | *Zea mays* | Iowa, USA | 2006 | NZ_CP033722 | Complete | 3.07 | - | PacBio | 236.0x | HGAP v. Can v. 1.5 |
| *C. m.* subsp. *nebraskensis* | *C. nebraskensis* | CFBP 7577 ϕ | *Zea mays* | ? | 2011 | QWED00000000 | Contig | 2.98 | 1272 | Illumina HiSeq | 6.0x | Velvet v. 1.2.10; SOAPdenovo v. 2.04; SOAPGapCloser v. 1.12 |
| *C. m.* subsp. *nebraskensis* | *C. nebraskensis* | DOAB 395 | *Zea mays* | Manitoba, Canada | 2014 | LSOE00000000 | Contig | 3.08 | 25 | Illumina | 110x | ABySS v. 1.5.2 |
| *C. m.* subsp. *nebraskensis* | *C. nebraskensis* | DOAB 397 | *Zea mays* | Manitoba, Canada | 2014 | LAKL00000000 | Contig | 3.06 | 28 | Illumina MiSeq | 89.0x | ABySS v. 1.5.2 |
| *C. m.* subsp. *sepedonicus* | *C. sepedonicus* | ATCC 33113$^T$ | *Solanum tuberosum* | Canada | 1980 | NC_010407 | Complete | 3.40 | - | Sanger AB 3700 | 8x | PHRAP, Gap4 |
| *C. m.* subsp. *sepedonicus* | *C. sepedonicus* | CFIA-Cs3N | *Solanum tuberosum* | Canada | ? | MZMM00000000 | Contig | 3.36 | 5 | PacBio | 536.0x | Celera Assembler v. 1 |
| *C. m.* subsp. *sepedonicus* | *C. sepedonicus* | CFIA-CsR14 | *Solanum tuberosum* | Canada | ? | MZMN00000000 | Contig | 3.41 | 6 | PacBio | 537.0x | Celera Assembler v. 1 |
| *C. m.* subsp. *tessellarius* | *C. tessellarius* | ATCC 33566$^T$ | *Triticum aestivum* | Nebraska, USA | 1978 | CP040788-CP040791 | Complete | 3.37 | - | PacBio RSII | 574.0x | HGAP v.4 |
| *C. michiganensis* | *C. tessellarius* | CFBP 3399 ¥ | *Tulipa* sp. | Netherlands | 1987 | JAHEWV000000000 | Contig | 3.26 | 68 | Illumina NovaSeq | 257.0x | SPAdes v. 3.15.2 |
| *C. m.* subsp. *tessellarius* | *Clavibacter* sp. | DOAB 609 ¥ | *Triticum aestivum* (leaves) | Nebraska, USA | 1976 | LQXA00000000 | Contig | 3.30 | 61 | Illumina MiSeq | 380.0x | ABySS v. 1.5.2 |
| *C. michiganensis* | *Clavibacter* sp. | CFBP 8017 ¥ | *Solanum lycopersicum* (seeds) | Netherlands | 2006 | MDJY00000000 | Contig | 3.17 | 65 | Illumina HiSeq | 90.0x | SPAdes v. 3.6 |
| *C. zhangzhiyongii* | *C. zhangzhiyongii* | DM1 | *Hordeum vulgare* (seeds) | Australia | 2017 | NZ_CP061274 | Complete | 3.10 | - | PacBio; Illumina | 450.0x | SMRT Link v. v5.1.0; Arrow v. v2.3.3 |
| *C. zhangzhiyongii* | *C. zhangzhiyongii* | DM3 | *Hordeum vulgare* (seeds) | Australia | 2017 | JAFEUE000000000 | Contig | 3.02 | 684 | Illumina HiSeq | 219.8x | Velvet v. 1.2.10 |
| *Rathayibacter toxicus* | *R. toxicus* | WAC3373 ¶ ǂ | *Phalaris paradoxa* | Australia | 1978 | NZ_CP013292 | Complete | 2.35 | - | PacBio RSII | 99x | HGAP |

| | | | | | | | | | | | | |
|---|---|---|---|---|---|---|---|---|---|---|---|---|
| *R. iranicus* | *R. iranicus* | NCCPB 2253 ¶ | *Triticum aestivum* (seed heads) | Iran | 1961 | NZ_CP028130 | Complete | 3.44 | - | PacBio | 156.0x | Celera Assembler v. 2013 |

ᵀ Indicates type strain of the bacterial cultures.

¶ Strains selected as outgroups for the multi-locus sequencing analysis (MLSA).

ⱡ Strain selected as an outgroup for the average nucleotide identity (ANI) computation and ANI-based phylogeny test.

ϕ Strains excluded from the core-genome-based phylogenomic study.

¥ strains used for taxonomy delineation assessment based on the digital DNA-DNA hybridization (dDDH) calculations as implemented in the Type Strain Genome Sever (TYGS) platform.

- Non applicable.

? Unknown information.
